# *Pogz* deficiency leads to abnormal behavior, transcription dysregulation and impaired cerebellar physiology

**DOI:** 10.1101/437442

**Authors:** Reut Suliman, Ben Title, Yahel Cohen, Maayan Tal, Nitzan Tal, Bjorg Gudmundsdottir, Kristbjorn O. Gudmundsson, Jonathan R Keller, Guo-Jen Huang, Yosef Yarom, Sagiv Shifman

**Author notes:** Equal contribution (Reut Suliman, Ben Title). Correspondence: Yosef Yarom, Department of Neurobiology, The Institute of life sciences, The Hebrew University of Jerusalem, Edmond J. Safra campus, Jerusalem 91904, Israel. Sagiv Shifman, Department of Genetics, The Institute of Life Sciences, The Hebrew University of Jerusalem, Edmond J. Safra campus, Jerusalem 91904, Israel.

## Abstract

Genes implicated in autism spectrum disorder (ASD) are enriched with chromatin regulators, but the mechanisms leading to the abnormal behavior and cognition are still unclear. Animal models are crucial for studying the effects of mutations on brain function and behavior. We generated conditional knockout mice with brain-specific mutation in *Pogz*, a heterochromatin regulator recurrently mutated in ASD and other neurodevelopmental disorders, and demonstrated that these mice display phenotypes that resemble the human condition. *Pogz* deficiency led to smaller brain, growth impairment, motor learning deficits, and increased social interactions that mimic the human overly friendly phenotype. At the molecular level, reporter assay indicated that POGZ functions as a negative regulator of transcription through its interaction with HP1 proteins. In accordance, we found a significant upregulation of gene expression, most notably in the cerebellum. Furthermore, *Pogz* deficiency was associated with a significant reduction in the firing frequency of simple and complex spikes in cerebellar Purkinje cells with no changes in their intrinsic properties. Overall, our findings support a mechanism linking heterochromatin dysregulation to cerebellar circuit dysfunction and to motor and social abnormalities in ASD.

## Introduction

The etiology of autism spectrum disorder (ASD) has puzzled medical researchers for several decades, but in recent years, there have been several breakthroughs. One of these is the increased appreciation regarding the importance of *de novo* mutations in ASD^1^. Exome sequencing studies revealed that hundreds of genes may contribute to ASD when disrupted by *de novo* mutations^2–5^. Furthermore, many of the genes implicated in ASD were also found to be associated with other types of neurodevelopmental disorders (NDDs)^6–10^.

A major goal in ASD research is to identify common mechanisms that are shared by the numerous ASD-associated genes, aiming to explain why mutations in all of these different genes commonly lead to ASD. Systematic analyses of genes with *de novo* mutations in ASD showed enrichment for transcription regulators and chromatin modifiers^5,11^, strongly suggesting that alteration in transcription and chromatin organization might have key pathogenic roles in ASD. Specifically, we noted that several of the high-confidence ASD risk genes are involved in heterochromatin formation or participate in complexes that bind different isotypes of the human heterochromatin protein 1 (HP1) (e.g. *SUV420H1*^12^, *ADNP*^13^ and *POGZ*^14–18^). A key role of HP1 proteins is transcriptional silencing and modulation of chromatin architecture, including transcriptional repression of euchromatic genes^19^. Transcriptional repression is crucial for neurogenesis and function of post-mitotic neurons, whereas aberrant silencing can lead to NDDs^20,21^.To study the mechanisms by which mutations in chromatin-related genes contribute to ASD, we focused on the *Pogz* gene (POGO transposable element with ZNF domain). Not only is *Pogz* one of the most significantly associated genes with ASD, it is also consistently found to be a strong interactor with the three isotypes of HP1^22–24^. Recent studies provided compelling evidence that loss-of-function (LoF) mutations in *POGZ* are associated with abnormal development and behavior^14–18^. Many of the individuals with *POGZ* mutations have developmental delay, and more than half are diagnosed with ASD. Other phenotypes are intellectual disability, microcephaly, overly friendly behavior, short stature, hyperactivity and vision problems^16–18^. The disorder caused by *POGZ* mutations is now known as White-Sutton syndrome [MIM: 616364]. Despite the detailed phenotypes, the molecular, cellular and physiological mechanisms of this syndrome are still unclear.

To study the function of POGZ, we generated a mouse model with a heterozygous or homozygous brain specific deletion of the *Pogz* gene. We found additive effects of *Pogz* on growth, behavior, brain physiology and gene expression. Our study suggests that mutations in *Pogz* causes transcription derepression, which leads to circuit and neuronal dysfunction, particularly in the cerebellum, and eventually contribute to the behavioral and cognitive symptoms seen in humans with *POGZ* mutations.

## Results

### Brain-specific deletion of Pogz does not result in gross defects in brain anatomy

We studied how *Pogz* dosage affect phenotypic outcome by generating heterozygote and homozygote knockout (KO) of *Pogz*. Since a complete KO of *Pogz* cause early embryonic lethality ^25^, we crossed a conditional *Pogz* flox/flox (*Pogz*^fl/fl^) mice with transgenic Nestin^CRE^ (Nes^Cre/+^; *Pogz*^fl/+^) in order to produce heterozygous (Nes^Cre/+^; *Pogz*^fl/+^) and homozygous (Nes^Cre/+^; *Pogz*^fl/fl^) mutation restricted to the brain (hereafter referred to as *Pogz* cKO^+/−^ and *Pogz* cKO^−/−^) (Figure 1A). Littermates without Nestin^CRE^ were used as a control. Along the study, we included both males and female mice, and the data was analyzed with sex as a covariance. The efficient deletion of POGZ in the brain was validated by western blot and immunostaining (Figure 1B-C). The immunostaining with POGZ antibodies also showed widespread expression of POGZ in neurons across the postnatal and adult brain of control mice (Figure 1C, S1). To study whether *Pogz* deficiency causes any anatomical abnormalities, we used Nissl staining, which revealed no gross brain anatomical defects in *Pogz* cKO^−/−^ mice (Figure 1D). Immunostaining with different neural markers in early infant (P11) and adult mice did not show any detectable differences between *Pogz* cKO^−/−^ and control in the cerebellum, hippocampus and cortex (Figure S2), including the thickness of cortical layers (all *P* > 0.05) (Figure 1E-F). Furthermore, Golgi staining images revealed similar levels of dendritic spine density in the dentate gyrus of *Pogz* cKO^−/−^ and control (*P* = 0.98) (Figure S3).

**Figure 1.**
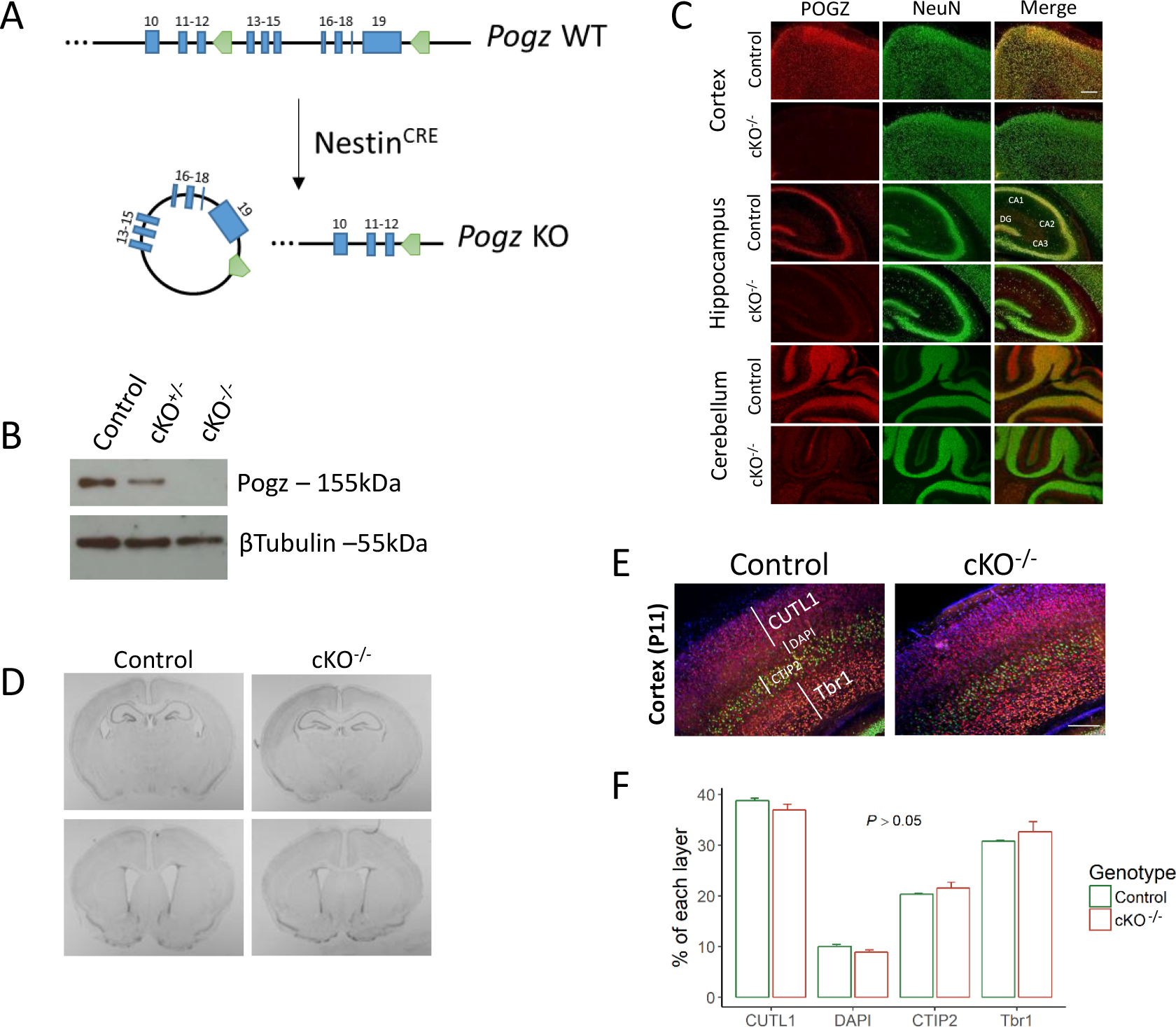
*Pogz* deficiency mouse model do not exhibit gross defects in brain anatomy. (A) Schematic overview of *Pogz* knockout strategy. *Pogz* gene exons 13-19 (blue) were bounded by *loxP* sites (green). *Pogz* cKO^+/−^ and *Pogz* cKO^−/−^ mice were generated with a mutation restricted to the brain, by crossing floxed *Pogz* mice with *Nestin* Cre mice. (B) Western blots analysis of whole cell lysates extracted from E14.5 mice cortices. (C) Immunofluorescence staining of *Pogz* cKO^−/−^ and control mice (P11) brains (cortex, hippocampus, cerebellum) using antibodies against POGZ (red) and NeuN (green). n = 3 controls, 3 cKO^−/−^. Scale bar = 100μm. (D) Nissl staining of coronal sections from *Pogz* cKO^−/−^ and control mice. (E) Immunostaining for specific cortical layers with known markers (CUTL1, CTIP2, Tbr1) and DAPI. Scale bar = 100μm. (F) Quantification of the relative size of each cortical layer was based on the length of layers stained with the markers or stained only by DAPI. n= 3 controls, 3 cKO^−/−^; all *P* > 0.05, Two-tailed t-test. Quantitative data are mean ± standard error of the mean (SEM).

### Pogz-deficient mice show growth delay, smaller absolute brain and decreased adult neurogenesis

We observed a significant reduction in the body size of *Pogz* cKO^−/−^ relative to control littermates, with heterozygotes (*Pogz* cKO^+/−^) showing an intermediate phenotype across development (*P* = 1.1×10^−12^) (Figure 2A-B). There was also a significant gene-sex interaction for body weight, with stronger effect in males (*P* = 0.016). As some individuals with POGZ mutations show microcephaly, we also measured brain weight at P11, and found a significantly lower absolute brain weight in *Pogz* cKO^−/−^ mice (*P* = 0.025) (Figure 2C). However, brain weight relative to body weight was significantly higher in *Pogz* cKO^−/−^ mice (*P* = 0.0062) (Figure 2D). Given the absolute smaller brain, the known role of POGZ in mitosis^24^ and the association of *POGZ* with microcephaly in humans, we hypothesized that POGZ may affect neurogenesis. We studied how *Pogz* dosage influence adult neurogenesis in the dentate gyrus using the immature neurons marker, doublecortin (DCX) (Figure 2E). We found a linear decrease in neurogenesis as a function of *Pogz*-deficiency levels. On average, each *Pogz* mutated allele decreased neurogenesis by 15.5% (*P* = 0.00033) (Figure 2F). Lower levels of immature neurons can be caused by decreased proliferation and/or reduced cell survival. Our experiments showed that neurogenesis was decreased in *Pogz*-deficient mice due to reduced survival, as indicated by the number of BrdU+/NeuN+ labeled cells (21.3% reduction per *Pogz* mutated allele, *P* = 0.0044) (Figure 2G). In contrast, proliferation was not significantly associated with *Pogz*-deficiency levels. The number of cells labeled by the KI67 marker was not significantly associated with *Pogz*-deficiency levels (*P* = 0.099), but were significantly elevated in *Pogz* cKO^+/−^ mice relative to control (adjusted *P* = 0.033) (Figure 2H).

**Figure 2.**
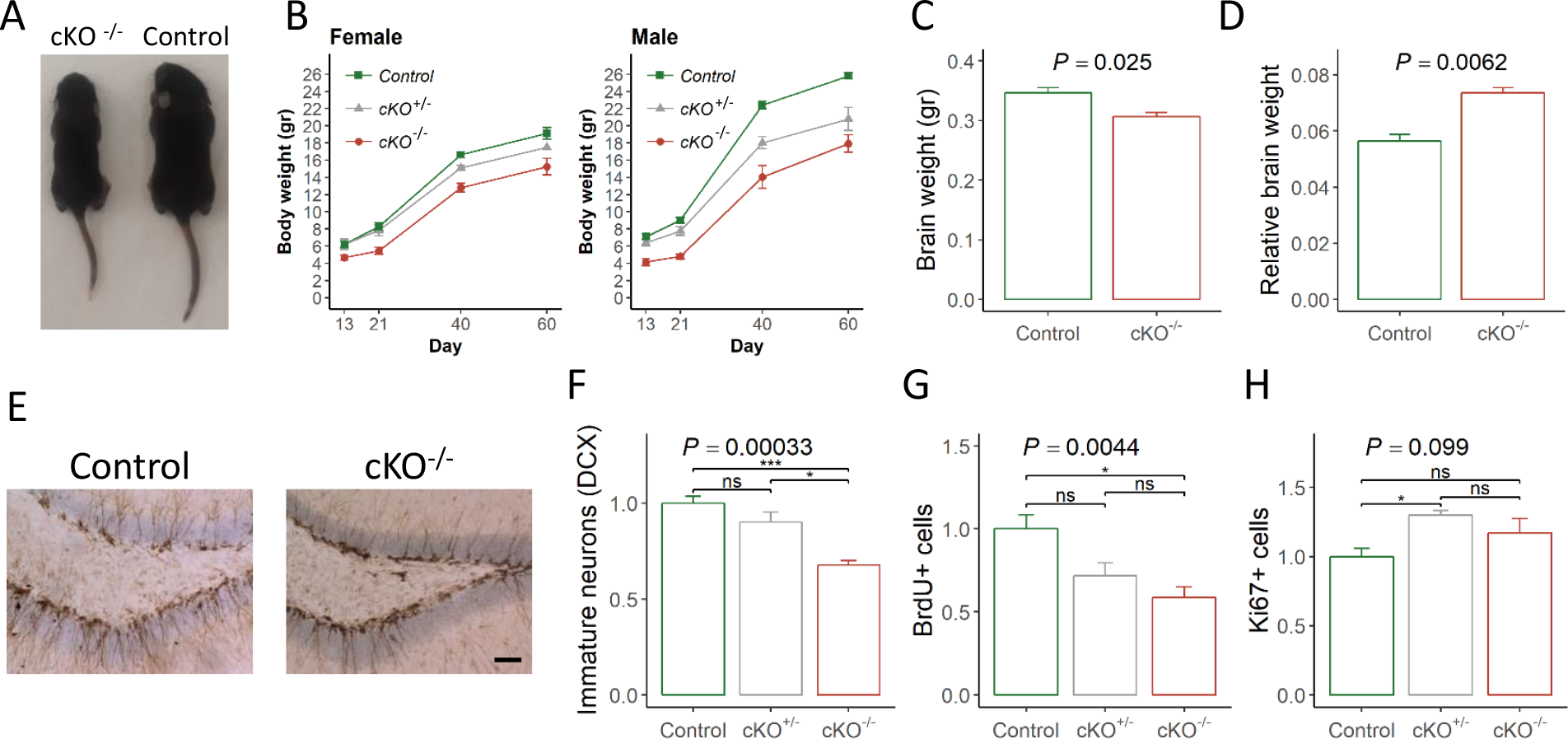
*Pogz*-deficient mice show growth delay, changes in brain size, and reduced neurogenesis. (A) Comparison of the body size of *Pogz* cKO^−/−^ and control mice littermates (P11) (B) Mice weight in four time points (P13, P21, P40, P60). Repeated measures ANOVA; n = 16 control, 7 cKO^+/−^, 7 cKO^−/−^, *P* = 1.1 × 10^−15^. (C) Absolute and (D) relative brain weight of control and *Pogz* cKO^−/−^ mice (P11). n = 4 control, 3 cKO^−/−^, Two-tailed t-test. (E) Immunostaining for Doublecortin (DCX) (positive DAB color) in the dentate gyrus (DG) of control and *Pogz* cKO^−/−^ mice. Scale bar = 50μm. Quantification of either (F) DCX positive cells (G) BrdU positive cells (H) Ki67 positive cells in the DG of control and *Pogz*-deficient mice. Numbers are proportion relative to control mice. n = 6 control, 3 cKO^+/−^, 3 cKO^−/−^. The *P*-values at the top of the plots are for association between the quantitative measurements and the number of intact *Pogz* alleles calculated with a linear regression model (unless stated otherwise). Pairwise comparisons between genotypes was calculated by Tukey HSD test and the significance is represented by: *, *P* < 0.05; ***, *P* < 0.001; ns, not significant. Quantitative data are mean ± SEM.

### Pogz-deficient mice showed abnormal motor and social behavior

One of the major phenotypes of White-Sutton syndrome is delayed psychomotor development ^14,18^. In addition, autistic features are common, with some individuals showing an overly friendly behavior. A summary of the tests performed relevant to the human phenotypes and the results obtained are presented in Table S1.

To test motor coordination and strength we used the accelerating rotarod and horizontal bar tests. In the rotarod. We found differences in motor learning between genotypes, as indicated by a significant trial by gene interaction (*P* = 0.023) and a significant improvement in performance (time on the rotating rod) only in control mice (*P* = 0.0015), but not in *Pogz* cKO^+/−^ (*P* = 0.094) or *Pogz* cKO^−/−^ mice (*P* = 0.57) (Figure 3A). Motor deficits were also evident in the horizontal bar test. The degree of *Pogz*-deficiency was significantly associated with reduced scores (*P* = 0.0022) (Figure 3B). A significant lower score was found for *Pogz* cKO^−/−^ compared to *Pogz* cKO^+/−^ (adjusted *P* = 0.0077) and control littermates (adjusted *P* = 0.0029). Motor deficits can affect locomotor activity, but the distance moved in the open field was not significantly altered by *Pogz* deficiency levels (*P* = 0.26) (Figure 3C).

**Figure 3.**
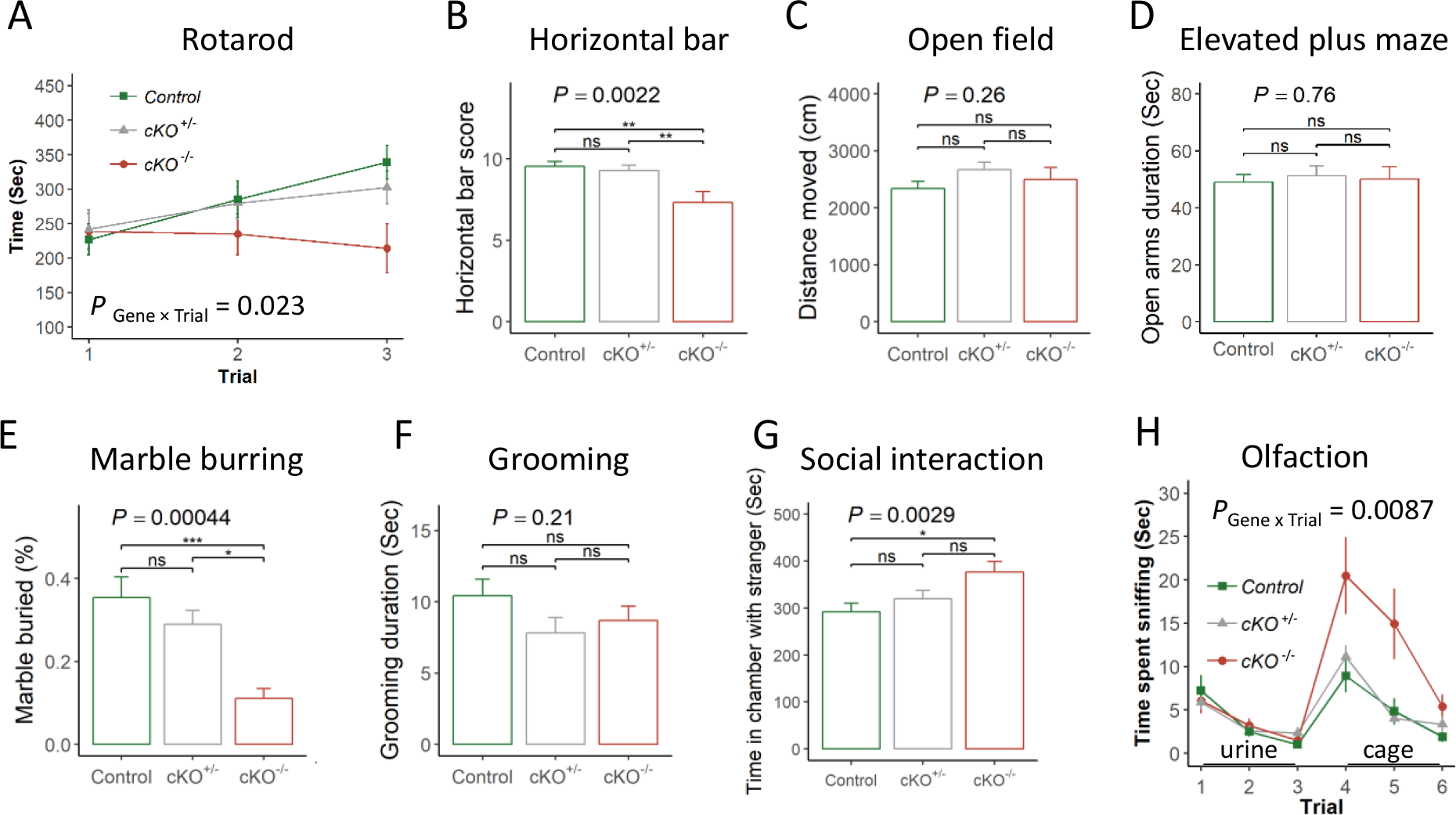
*Pogz*-deficient mice show abnormal motor and social behavior. (A) Time spent on the accelerated rotarod for three consecutive trials. Repeated measures ANOVA; n= 26 control, 25 cKO^+/−^, 11 cKO^−/−^. (B) Time taken to cross or fall from the horizontal bar calculated as a score. n= 13 control, 14 cKO^+/−^, 9 cKO^−/−^. (C) Total distance moved in the open field. n= 30 control, 28 cKO^+/−^, 11 cKO^−/−^. (D) Time spent in the open arms of the elevated plus maze. n= 40 control, 38 cKO^+/−^, 18 cKO^−/−^. (E) Percentage of marbles buried in the marble burying test. n= 13 control, 14 cKO^+/−^, 9 cKO^−/−^. (F) Time spent self-grooming during the open field. n= 30 control, 28 cKO^+/−^, 11 cKO^−/−^. (G) Time spent in chamber with a stranger mouse as measured in the three-chamber sociability test. n= 14 control, 14 cKO^+/−^, 11 cKO^−/−^. (H) Time spent sniffing urine and cage odors in the olfactory habituation – dishabituation test. Repeated measures ANOVA; n= 20 control, 23 cKO^+/−^, 9 cKO^−/−^. The *P*-values at the top of the plots are for association between the quantitative measurements and the number of intact *Pogz* alleles calculated with a linear regression model (unless stated otherwise). Pairwise comparisons between genotypes was calculated by Tukey HSD test and the significance is represented by: *, *P* < 0.05; ** *P* < 0.01; ***, *P* < 0.001; ns, not significant. Quantitative data are mean ± SEM.

To assess anxiety related responses, we performed the elevated plus maze and observed no significant differences in the time spent in the open arms (*P* = 0.76) (Figure 3D). Similarly, there was no significant difference in the duration of time spent in the center of the open field (*P* = 0.76) (Figure S4A). Next, we used the marble burying test to asses repetitive digging, which is also known to be influenced by antidepressants ^26,27^, and self-grooming as another measure of repetitive behavior. Marble burying was significantly reduced in *Pogz*-deficient mice (*P* = 0.00044) (Figure 3E), while grooming was not significantly different (*P* = 0.21) (Figure 3F).

We next analyzed social behavior in a three-chamber social interaction test. Consistent with the overly friendly phenotype seen in humans with *POGZ* mutations, *Pogz*-deficiency levels were significantly associated with increased duration of time in the chamber and near the cup with a stranger mouse (*P*_Chamber_ = 0.0029, *P*_Cup_ = 0.0071) (Figure 3G, S4B-C). In the social novelty test, there was no significant difference between genotypes (*P* > 0.3, Figure S4D-E). To test if *Pogz*-deficient mice can detect and differentiate different social odors, we tested the mice in an olfactory habituation - dishabituation task with urine odor following by cage swipes odor (from age and sex matched mice). All genotypes showed habituation to the odor and dishabituation to the novel odor, but there was a significant differences in sniffing time (genotype effect, *P* = 0.0022; gene by trial interaction, *P* = 0.0087). Consistent with increased social interest of *Pogz*-deficient mice, the sniffing time of cage swipes was significantly increased in *Pogz* cKO^−/−^ (*P* <0.05) (Figure 3H) but not for urine. In a buried food olfaction test there were no significant differences (*P* = 0.097) (Figure S4F).

### POGZ repress transcription via HP1

Since POGZ was found to be integral part of the HP1 protein complexes ^22–24^, we hypothesized that POGZ plays a key role in transcription regulation. Based on immunostaining, POGZ is distributed at pericentric heterochromatin (Figure 4A), in a similar way as HP1α and HP1β ^28,29^. To assess whether POGZ regulates transcription, we performed a dual-luciferase reporter assay in HEK293 cells. We transiently introduced a GAL4 DNA-binding domain (GAL4-DBD) fused with full-length POGZ (POGZ-FL). Transcription of the reporter gene was significantly inhibited by POGZ-FL (*P* = 9 × 10^−5^) (Figure 4B). POGZ interacts with HP1 proteins through a HP1-binding zinc finger-like (HPZ) domain (Figure 4C), rather than the canonical PxVxL motif ^24^. To test if the interaction with HP1 is essential for transcription repression we used POGZ with a mutation in the HPZ domain (POGZ-H840A) that affects HP1-binding ^24^. POGZ-H804H failed to repress reporter gene transcription (*P* = 0.4) (Figure 4B), suggesting that POGZ functions as a negative regulator of transcription through its interaction with HP1 proteins.

**Figure 4.**
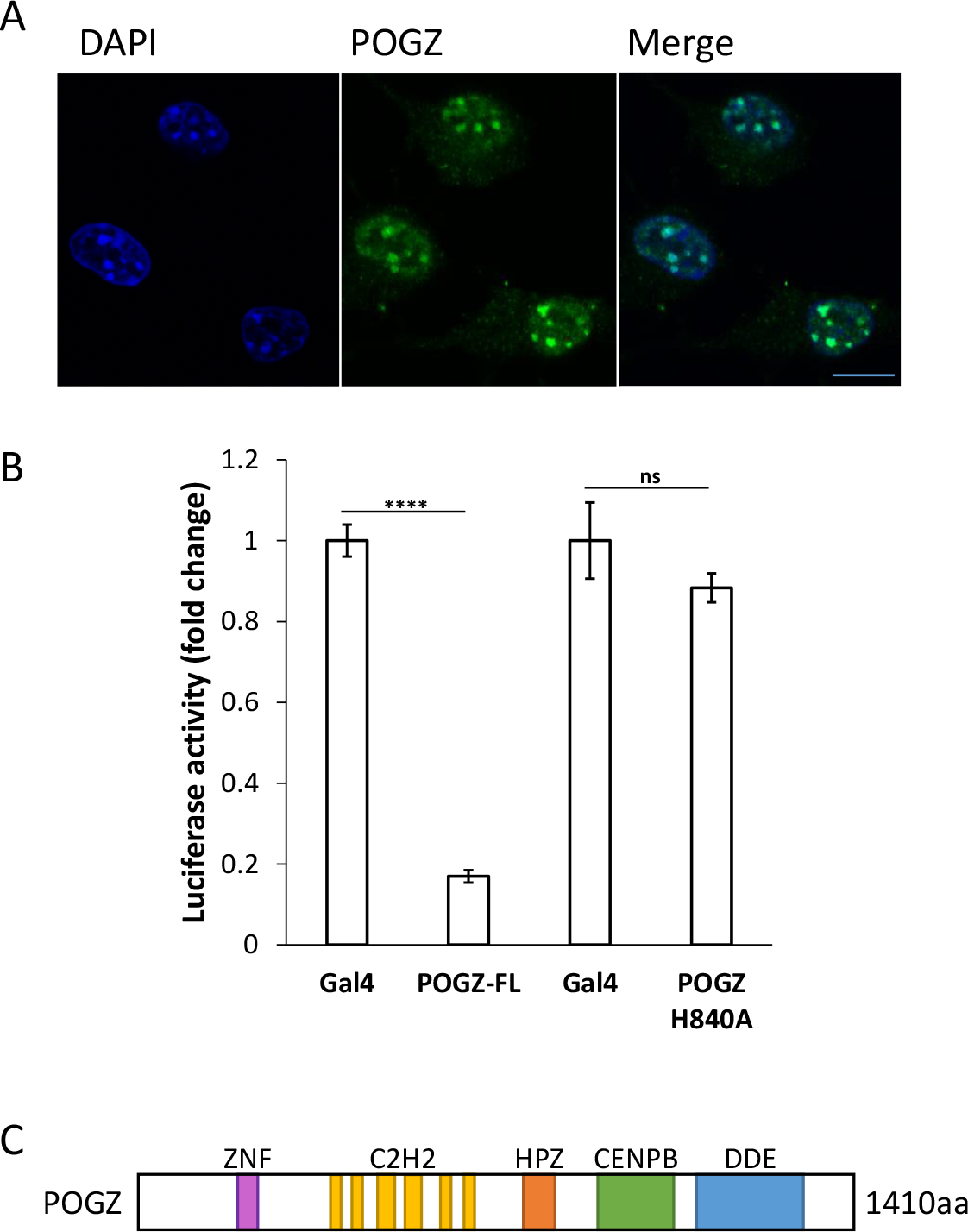
POGZ localizes to pericentric chromatin and repress transcription via HP1 proteins. (A) Immunofluorescence staining of mouse neuronal cells (N2A) using antibody against POGZ (green) and DAPI (blue). (B) Fold change in luciferase activity in HEK293 cells expressing: Gal4, Gal4 DNA binding domain (DBD) fused to full-length POGZ (POGZ-FL), or Gal4-DBD fused to POGZ with a mutation in the HPZ sequence that changes the conserved amino acid Histidine to Alanine (POGZ H840A) and affects HP1-binding. Two-tailed t-test; ns, not significant. Data are mean ± SEM. (C) POGZ protein structure including the C2H2 zinc-finger motifs (purple and yellow), HPZ domain (orange), CENP-B DNA-binding domain (green) and DDE domain (blue). The amino-acid sequence encoding the HPZ domain (amino-acid residues 791-850) is essential for the interaction with HP1 proteins.

### Transcriptional dysregulation in the brain of Pogz-deficient mice

Given that POGZ is involved in transcriptional repression, we wanted to study how POGZ affect gene expression in the brain. While *Pogz* has widespread expression in the mouse brain, it is highly expressed in the cerebellum, in both human and mouse (Figure 5A-B). We therefore focused on gene expression in the cerebellum of adult mice and compared it to the hippocampus, where *Pogz* is less expressed. We performed the differential expression analysis using RNA-seq on six RNA samples from each genotype, *Pogz* cKO^−/−^, *Pogz* cKO^+/−^ and control littermates, including males and females. To identify differential expression that is associated with *Pogz* dosage, we used a statistical model that accounted for sex (gene expression as a linear function of the number of intact *Pogz* alleles), across genes that were robustly expressed (counts per million (cpm) > 1 in at least a third of the samples; hippocampus: n = 14,339, cerebellum: n = 14,027). At significance cutoffs corresponding to false discovery rate (FDR) < 0.05, we found 636 genes that were differentially expressed in the hippocampus, among them 46 with fold change (average effect of allele substitution) above 1.5 and 15 with fold change > 2 (Figure 5C-D; Table S2). In the cerebellum, we identified a threefold increase in the number of differentially expressed genes: 1916 at significance cutoffs corresponding to FDR < 0.05, 196 with fold change above 1.5 and 52 with fold change > 2 (Figure E-F; Table S2). Consistent with role of POGZ in gene repression, the majority of differentially expressed genes with absolute fold change above 1.5 (hippocampus: 67%, *P* = 0.026; cerebellum: 77%, *P* = 4.6 × 10^−14^) or 2.0 (hippocampus, 93%, *P* = 0.00098; cerebellum: 83%, *P* = 2.0 × 10^−6^) were upregulated. There was no significant difference in the proportions when genes with absolute fold change below 1.5 were included (hippocampus: 47%, *P* = 0.14; cerebellum, 52%, *P* = 0.079).

**Figure 5.**
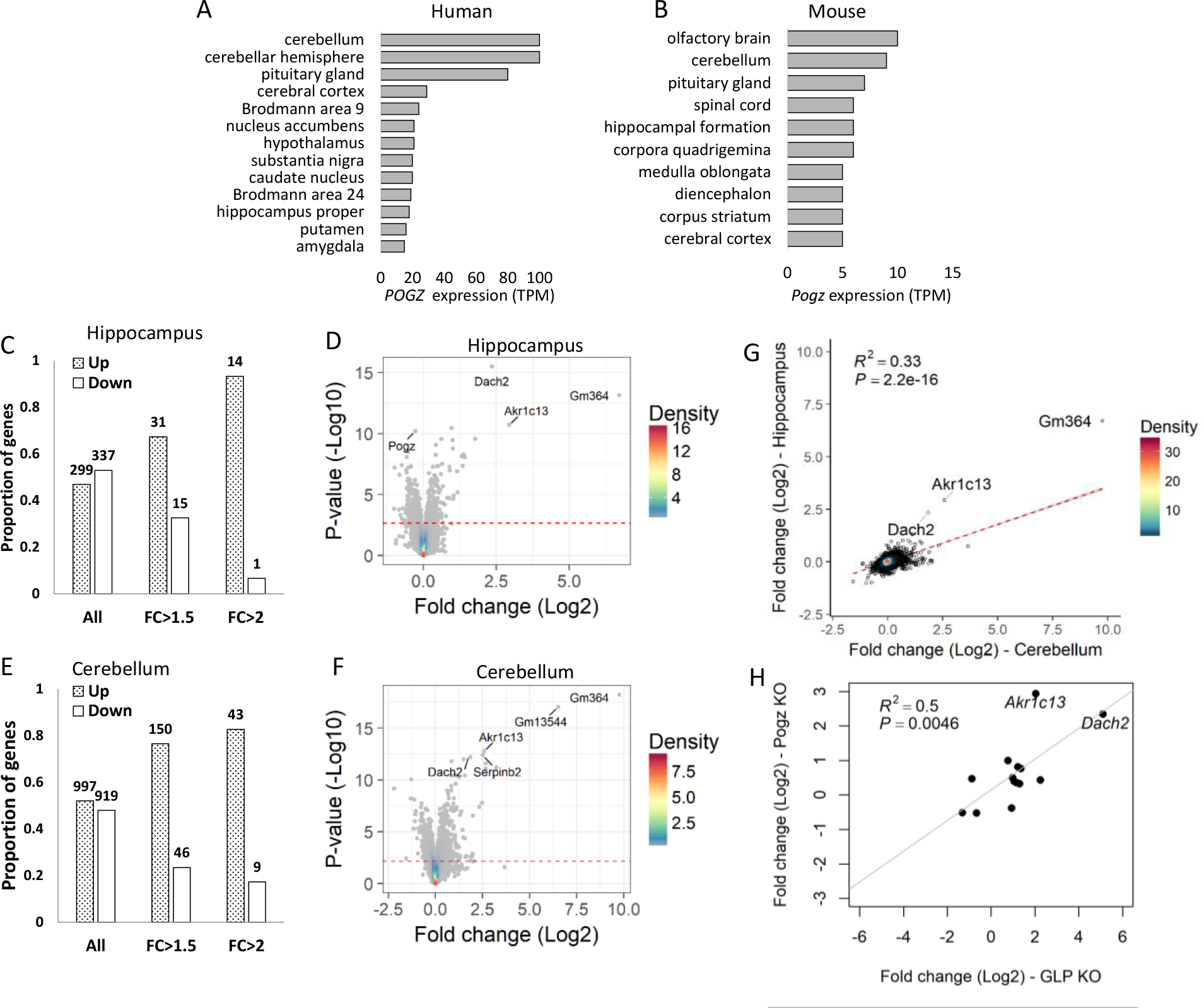
*Pogz* deficiency leads to transcriptional dysregulation. (A) Levels of *POGZ* expression in different brain areas of human and (B) *Pogz* expression in mouse (data from the FANTOM5 dataset). Values are transcripts per million (TPM) (C, E) Proportion of up- and down- regulated genes in the (C) hippocampus or (E) cerebellum of *Pogz*-deficient mice. Bars correspond to all genes (left), genes with fold change above 1.5 (middle) and genes with fold change above 2 (right) (D, F) Volcano plot showing differentially expressed genes in the (D) hippocampus or (F) cerebellum. The red line indicates FDR = 0.05. (G) Correlation between fold changes of gene expression in the hippocampus and the cerebellum (H) Correlation between fold changes of significantly differentially expressed genes in the hippocampus of *Pogz*-deficient mice and GLP KO mice ^36^.

Despite the differences in the number of differentially expressed genes between the hippocampus and cerebellum, the changes in expression were related. First, 41.1% of genes differentially expressed in the hippocampus were also differentially expressed in the cerebellum (*P* < 2.2 × 10^−16^). Second, the fold change across all commonly expressed genes was significantly correlated between the hippocampus and cerebellum (R^2^ = 0.33, *P* < 2.2 × 10^−16^), but the slope of the regression line indicted that the gene expression changes in the hippocampus are consistently lower compared to the cerebellum (Figure 5G). Three genes were upregulated in the hippocampus more than expected based on the fold change in the cerebellum (*Dach2*, *Akr1c13*, and *Gm364*). *Akr1c13* is a rodent-specific gene expressed exclusively in stomach, liver and ileum ^30^; *Gm364* is a mouse-specific gene, highly expressed in the adult mouse testis. *Dach2 is* a transcription factor ^31^ that is the most significant differentially expressed gene in the hippocampus (adjusted *P* = 4.5 × 10^−12^), and one of the top significant genes in the cerebellum (adjusted *P* = 4.9×10^−9^). Relative to control littermates, *Dach2* was upregulated in the hippocampus of *Pogz* cKO^−/−^ mice by 24.3 fold and 4.5 fold in *Pogz* cKO^+/−^ (Figure S5A). In the cerebellum, *Dach2* was upregulated by 12.5 fold in *Pogz* cKO^−/−^ mice and by 3.8 fold in *Pogz* cKO^+/−^ mice (Figure S5B). We confirmed the upregulation of *Dach2* by quantitative PCR (qPCR) in the hippocampus, cerebellum, and the cortex (Figure S4C).

*Dach2* was previously identified as a transcriptional repressor that induces organogenesis and myogenesis ^32,33^. We therefore hypothesized that while upregulated genes may be enriched for *Pogz* targets, downregulated genes may be enriched with targets of *Dach2*. To test this, we checked downregulated and upregulated genes for the enrichment of histone marks using the *Enrichr* software ^34,35^ and the histone modification gene-set library (ENCODE Histone Modifications 2015). Consistent with our hypothesis and the role of *Dach2* in myogenesis, genes downregulated in hippocampus or cerebellum (FDR < 0.05 & fold change < −1.5) were most significantly enriched for the repressive histone mark H3K27me3 in mouse myocyte (adjusted *P* = 1.2 × 10^−6^) (Figure S6A), while upregulated genes (FDR < 0.05 & fold change > 1.5) showed the strongest enrichment for H3K27me3 in mouse cerebellum (adjusted *P* = 1.2 × 10^−8^) (Figure S6B).

Among the downregulated genes marked by H3K27me3 in mouse myocyte, the growth hormone secretagogue receptor (*Ghsr*) may be associated with the delayed growth of *Pogz*-deficient mice (downregulated in the hippocampus by 3.1 fold in *Pogz* cKO^−/−^ mice; adjusted *P* = 0.00011) (Table S3).

In a previous study, conditional knockout in forebrain postnatal neurons of GLP/G9a resulted in derepression of non-neuronal genes in multiple brain regions, including the hippocampus (the cerebellum was not tested) ^36^. G9a and GLP (*Ehmt1/Ehmt2*) are two genes in the histone methyltransferase complex that interacts with HP1 and POGZ, are responsible for mono- and di methylation of H3K9, and are required for transcription repression at euchromatic and facultative heterochromatin ^37–39^. To test if POGZ control similar set of genes we compared the differential gene expression in the hippocampus between the two studies. Remarkably, we found a highly significant overlap between the set of genes we identified to be differentially expressed in *Pogz*-deficient mice and the sets of differentially expressed genes in GLP cKO (34.1%; *P* = 6.3 × 10^−17^) or G9a cKO (21.7%; *P* = 2 × 10^−14^). In both GLP and G9a cKOs, the most significant differentially expressed gene was *Dach2*, but we also found a significant correlated changes in expression for the entire overlapping set of differential expressed genes between the mouse models (GLP vs. *Pogz*: r^2^ = 0.50, *P* = 0.0046; G9a vs. *Pogz*: r^2^ = 0.38, *P* = 0.014)(Figure 5H).

To identify biological Process affected by POGZ, we analyzed the differentially expressed genes for enrichment of gene ontology. The differentially expressed genes (FDR < 0.05) in the cerebellum were enriched for genes involved in Axon guidance (KEGG pathway, FDR = 2.3 × 10^− 4^), and skeletal system development (Biological processes, FDR = 2.0 × 10^−7^). In the hippocampus, the differentially expressed genes were enriched for genes involved in Ras signaling (KEGG pathway, FDR = 0.0031), Calcium signaling/calcium ion binding (KEGG pathway, FDR = 0.037), and transmembrane receptor protein tyrosine kinase activity (molecular function, FDR = 0.0014) (Table S4). To investigate whether changes in gene expression might underlie the observed phenotypes, we tested for mouse phenotypes (MGI mammalian phenotype 2017) associated with the differentially expressed genes. We found that genes differentially expressed in the cerebellum were most significantly enriched with abnormal joint morphology (FDR = 0.0033) and postnatal growth retardation (FDR = 0.045), while in the hippocampus the most significant enriched mouse phenotype was decreased body weight (FDR = 0.032) (Table S5).

### Pogz-deficient mice show reduction in spontaneous firing rate of Purkinje cells in both simple and complex spike

As the cerebellum is associated with the motor and transcriptional abnormalities of *Pogz*-deficient mice, we examined cerebellar activity in anesthetized animals, focusing on Purkinje cells (PCs) activity. To that end, we performed extracellular recordings of PCs in lobules V-VI (examples of unit recording are shown in Figure 6A). Complex spikes (CS) were readily identified and frequencies of spontaneous simple spike (SS) and CS were calculated. We observed that *Pogz* deficiency levels were significantly associated with a reduction in spontaneous SS frequency (*P* = 0.00012; Control: 45.0±28.8 Hz; cKO^+/−^: 32.5±18.0 Hz; cKO^−/−^: 20.4±16.0 Hz) (Figure 6B). Similarly, a significant reduction in spontaneous CS frequency was associated with *Pogz* deficiency levels (*P* = 0.00030; Control: 0.87±0.60 Hz; cKO^+/−^: 0.80±0.48 Hz; cKO^−/−^: 0.37±0.22 Hz) (Figure 6C).

**Figure 6.**
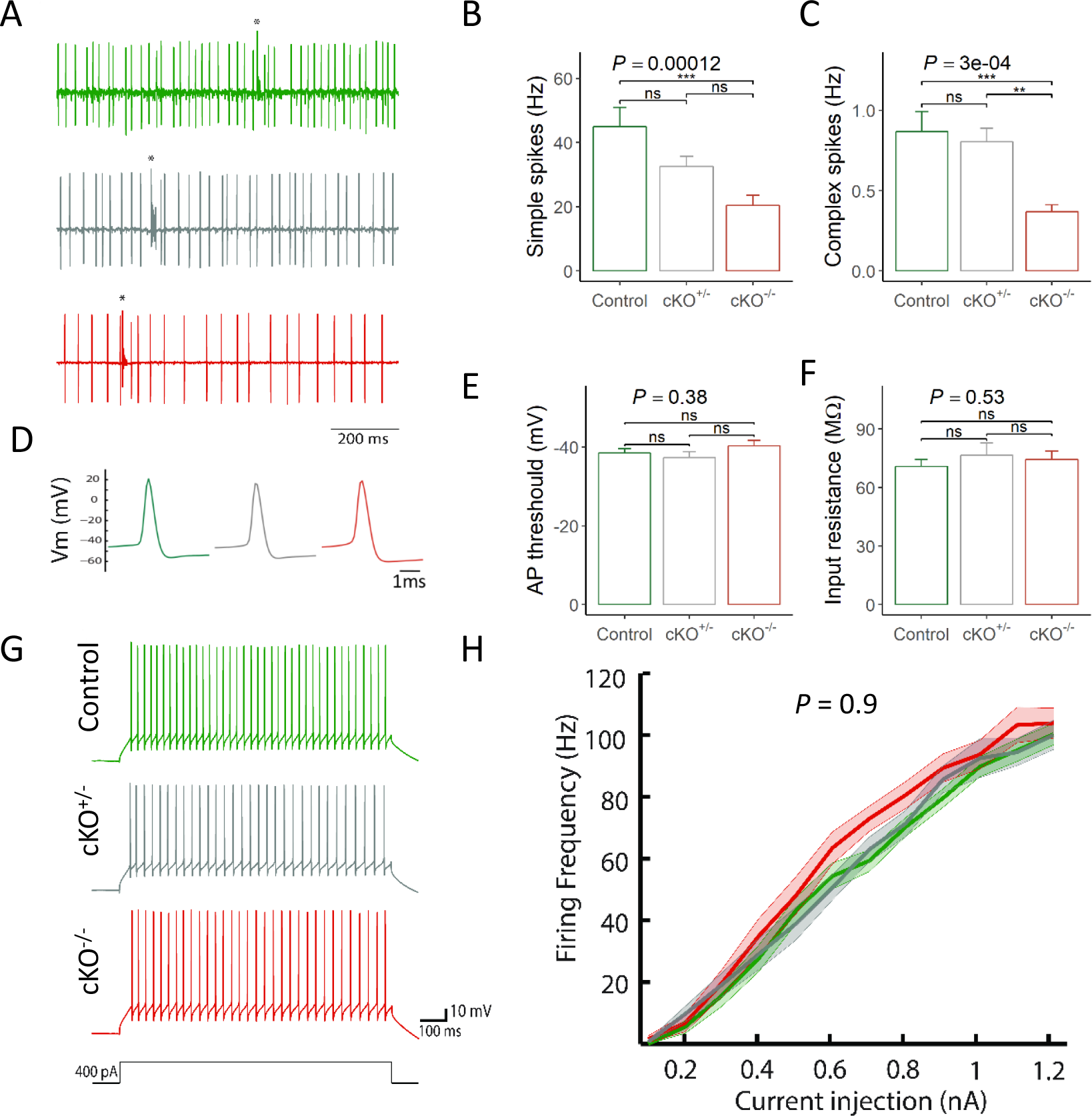
*Pogz*-deficient mice show reduced spontaneous firing frequency of Purkinje cells *in-vivo*, which cannot be explained by changes in their intrinsic properties. (A) Single unit activity recorded from PCs in lobules V-VI of the cerebellum. Asterisks indicate complex spikes. (B) Simple spikes and (C) Complex spikes spontaneous firing frequency. n {cells/animals}= 23/5 control, 32/5 cKO^+/−^, 25/3 cKO^−/−^ (D) Example traces of action potentials (AP) evoked by rheobase current injection. (E) AP threshold of PCs. n= 28/3 control, 21/2 cKO^+/−^, 21/2 cKO^−/−^ (F) Input resistance of PCs. n= 25/3 control, 15/2 cKO^+/−^, 21/2 cKO^−/−^ (G) Example traces of action potentials evoked by 400pA current injections. (H) The current-to-firing frequency (I-f curve). Repeated-measures ANOVA; n= 28/3 control, 21/2 cKO^+/−^, 21/2 cKO^−/−^. The *P*-values at the top of the plots are for association between the quantitative measurements and the number of intact *Pogz* alleles calculated with a linear regression model (unless stated otherwise). Pairwise comparisons between genotypes was calculated by Tukey HSD test and the significance is represented by: ** *P* < 0.01; ***, *P* < 0.001; ns, not significant. Quantitative data are mean ± SEM.

### Purkinje cells show no changes of intrinsic properties in Pogz-deficient mice

The reduction in CS frequency must be due to a change in the frequency of the olivary input^40^. However, a reduction in SS firing frequency can be either due to a change in the electrical properties of the PC or a change in the excitatory and inhibitory inputs into the PC. Therefore, we performed whole cell current-clamp recordings to investigate PC’s neuronal excitability. First, we examined the shape and threshold of the action potential (AP), using depolarizing current step. There was no significant difference in shape and AP threshold (*P* = 0.38; Control: −38.5±6.0 mV; cKO^+/−^: −40.3±6.6 mV; cKO^−/−^: −37.4±6.8 mV) (Figure 6D-E). Second, we measured PC input resistance using a short, low intensity, negative current injection (−50 pA of 100ms; see Materials and Methods). There was also no significant difference in PC input resistance (*P* = 0.53; Control: 70.8±17.8 MΩ; cKO^+/−^: 76.6±24.2 MΩ; cKO^−/−^: 74.4±20.1 MΩ) (Figure 6F). Next, we examined the responses for prolonged current injections to extract the current-frequency curve (I-f) (Figure 6G-H). All genotypes showed a similar I-f curve with no significant differences (*P* = 0.90). Finally, we examined the properties of the hyperpolarization-activated cationic current I_h_, which is known to play a major role in the electrophysiological properties of PCs with high relevance to the performance of motor learning behaviors^41–43^ and has been suggested as a target of another autism-associated *SHANK3* gene^71^. We found no differences between genotypes in either the voltage dependence of the I_h_ current (*P* = 0.79) (Figure S7A-B) or I_h_ current kinetics (Fast τ: *P* = 0.53; control: 139.9±33.9 ms; cKO^+/−^: 124.6±36.3 ms; cKO^−/−^: 150.8±54.0 ms; Slow τ: *P* = 0.07; control: 1.02±0.22 sec; cKO^+/−^: 0.88±0.30 sec; cKO^−/−^: 0.77±0.17 sec) (Figure S7C-D). These findings suggest that the observed reduction in SS firing frequency cannot be attributed to changes in PCs excitability.

## Discussion

We report the first characterization of *Pogz* knockout mouse in the brain, which revealed significant findings across genomic, cellular, physiological and behavioral dimensions of neurobiology. Since we studied mice that are both heterozygous and homozygous for the mutation in *Pogz*, we were able to show that the dosage of *Pogz* is required for repression of transcription in the brain, and is additively associated with the degree of abnormal behavior and physiology. Our results also suggest that POGZ is required for the proper function of the cerebellum, a brain region that has been linked consistently with ASD ^44,45^.

The phenotypes that we identified in the *Pogz*-deficient mice resemble several of the characteristics found in human individuals with *POGZ* mutations (White-Sutton syndrome)^14,17,18^. For example, the growth delay and small brain in mice may be related to the short stature and microcephaly seen in patients. The increased social interactions in mice may be related to the ASD diagnosis and overly friendly phenotype observed in human individuals with *POGZ* mutations. Furthermore, *Pogz*-deficient mice showed deficits in motor performance and learning, and previous study demonstrated that impaired motor skills in individuals with ASD are associated with *de novo* mutations across multiple genes ^46^. However, one should note that a complex genotype-phenotype relationship was reported for White-Sutton syndrome as well as for other syndromes associated with ASD, with a large phenotypic heterogeneity even between individuals that carry similar types of loss-of-function mutations. For example, microcephaly was reported only in small proportion of cases with White-Sutton syndrome, and a formal ASD diagnosis was reported for around half of the individuals ^14^. Therefore, it should not be expected that a mouse model would recapitulate all the abnormalities seen in patients.

We demonstrated that POGZ is able to repress transcription as a part of being in complex with HP1 proteins, and as consequence, mutations in *Pogz* lead to abnormal derepression of multiple genes in the brain. The overlap with genes differentially expressed in GLP or G9a mutant mice support the involvement of POGZ in a repression complex together with HP1 and GLP/G9a proteins. Indeed, previous studies showed the POGZ interact with HP1 and GLP/G9A proteins ^38^. Furthermore, mutations in GLP were reported in individuals with intellectual disability and ASD (Kleefstra Syndrome) ^36^. Thus, Kleefstra Syndrome and White-Sutton syndrome may be considered as related syndromes.

Multiple line of evidence point to the cerebellum as a critical brain region affected by *Pogz* deficiency. Not only that *POGZ* shows the highest expression in the human cerebellum relative to other brain regions, and this is similar in the mouse, we find abnormal motor performance and learning in *Pogz*-deficient mice, and reduced firing rates of cerebellar PCs. The reduction in SS activity can be due to alterations in PC intrinsic properties; however, basic properties such as input resistance or current frequency relationships were unaffected by *Pogz* deficiency. Thus, the reduction in SS activity should reflect a reduction in the mossy fibers input, reduction in the sensitivity of granule cells and/or changes in the local inhibitory network. Reduction in SS firing frequency of PC was previously described in a number of ASD mouse models but this was attributed to the change in PC properties ^47,48^. An increase in granule cells sensitivity has also been reported^49^, but it cannot account for our observed reduction SS activity. Finally, enhanced inhibitory input onto PC, which has been reported in *Shank2* model, can account for the reduction in SS activity^50^.

The reduction in CS activity is of special interest. First, to the best of our knowledge it is the first report on reduction of CS in ASD mouse model. Second, a parallel reduction in CS and SS is rather unusual given that these two types of PC activity are known to demonstrate reciprocal relationship. Namely, in any process that involve an increase in CS activity, a reduction in SS was always observed^51–53^. Therefore, a reduction in both types, suggests an overall reduction in cerebellar activity. Finally, a reduction in CS activity is bound to reflect a reduction in climbing fiber input. Given the proposed prominent role of climbing fiber in motor learning processes^54^, it can account for our observation on reduced motor learning.

In summary, our results support the role of mutations in *POGZ* as causing neurodevelopmental disorders, with symptoms resulting from transcription dysregulation and cerebellar circuit dysfunction.

## Materials and Methods

### Generation of brain specific Pogz-deficient mice

*Pogz* conditional knockout mice on the background of C57/BL6 were previously generated^25^ using the Cre-lox system, with loxP sites flanking exons 12 to 19. *E*xons 12 to 19 include the DDE transposas domain, the CENP DNA binding domain and a part of the zinc fingers domain that binds HP1 proteins (HPZ). To create a mutation that is restricted to the brain, we crossed the conditional *Pogz* mice with Nestin^CRE^ transgenic mice. All mice were tail genotyped using KAPA mouse genotyping kit (KAPA Biosystems KK7302(. PCR primers used were: 5’- AATTAAAGGCAGACCTAGCAGGTGGAGG- 3’ (forward) and 5’- TAGCACCGC AGACTGCTATCTATTCCTG (reverse) for loxP sequence, 5’- GCGGTCTGGCAGTAAAAACTATC- 3’ (forward) and 5’- GTGAAACAGCATTGCTGTCACTT -3’ (reverse) for Nestin-Cre. All mouse studies were approved by the Institutional Animal Care and Use Committees at The Hebrew University of Jerusalem.

### Western blot analysis

Proteins were extracted from cortices of E14.5 embryos for all three genotypes (control, *Pogz* cKO^+/−^ and *Pogz* cKO^−/−^) using modified RIPA buffer (50mM Tris pH 8.0, 150mM NaCl, 5mM EDTA pH 8.0, 1% Triton x100, 0.5% sodium deoxycholate, 0.1% SDS) mixed with protease inhibitors (Sigma-Aldrich). Each tissue sample was ruptured and agitated in RIPA for 1 hour in 4 degrees. The lysates were cleared by centrifugation (20 minutes; 12,000 rpm; 4 degrees). Before loading, Laemmli sample buffer X5 with 2% β-mercaptoethanol was added to the samples following by boiling at 95 °C for 5 minutes. Protein samples were loaded onto 8% acrylamide gel, and wet method was used to transfer the proteins to a PVDF membrane (Immobilon-P, Millipore IPVH00010). Membranes were air-dried for blocking followed by an overnight incubation in Primary antibodies. The next day, membranes were washed in PBST and incubated in secondary antibodies for 1 hour at RT. After another three PBST washes, immunoblots were visualized using SuperSignal chemiluminescent HRP substrates according to manufacturer’s instructions (Thermo Scientific).

### Quantitative PCR

cDNA was synthesized using the Superscript III First-Strand kit for qPCR (Invitrogen 18080093). Real-time PCR was performed using the SsoAdvanced™ Universal SYBR^®^ Green Supermix (BIO-RAD). Fluorescence was monitored and analyzed in a Bio-Rad C1000 Thermal Cycler with a CFX96 real-time system. All experiments were performed in triplicates and analyzed using the 2^(−ΔΔCT) method, using GAPDH as the reference gene. Primers: Pogz exons’ 17-19 junction: 5’- GTGATGTCATTGAGGAC-3’ (forward) and 5’-TAGTGTGACAAGAGTCTT-3’ (reverse); murine GAPDH: 5’-TGTTCCTACCCCCAATGTGT-3’ (forward) and 5’-ATTGTCATACCAGGAAATGAGCTT-3’ (reverse); murine Dach2: 5’-AGTCATGAAGTCACCCTTGGA-3’ (forward) and 5’- TTGGTCAACAGAGTCTCCACA-3’ (reverse); murine Ghsr: 5’-GCTGCTCACCGTGATGGTAT-3’ (forward) and 5’-ACCACAGCAAGCATCTTCACT-3’ (reverse).

### Luciferase reporter assay

HEK293 cells were cultured on 48-well cell culture plates overnight. Cells were then transiently transfected with 0.1 μg UAS-SV40-luc (luciferase reporter) and 0.04 μg pLR-TK (renilla internal control) and either of 0.25 μg pCDNA3.1-GAL4-POGZ-FL (expressing GAL4 fused to full length human POGZ) or 0.25 μg pCDNA3.1-GAL4-POGZ-H840A (expressing GAL4 fused to human POGZ mutated in the HPZ domain) using TransIT-2020. Relative luciferase activity was measured, in three independent experiments, after 48 h using the Dual-Luciferase Reporter system (Promega) according to the manufacturer’s protocols.

### Brain sectioning and staining

2-4-month-old mice were euthanized using Isoflurane USP (Piramal critical care, NDC 60307-110-25). The brain was then removed and fixed overnight in 4% PFA at 4 degrees. Brains were than incubated in 30% sucrose prior to sectioning. 40μm brain sections were made using Leica “SM2000R Sliding” microtome and stored in 2:1:1 PBS/Glycerol/Ethylene Glycol at −20 degrees. Mounted slides were incubated in 0.01M citrate buffer pH=6 and heated above 95⁰ for 15 minutes. The slides were then washed in PBS and incubated in 3% H_2_O_2_ for 10 minutes. Primary antibodies were diluted in 0.5% triton-PBS solution and 2% normal serum (host animal varies according to secondary antibody) and incubated overnight in a humid chamber in RT. For IHC staining: on the following day, slides were washed with PBS and incubated with secondary biotinylated antibody diluted in 0.5% triton-PBS and 2% for 1.5 hours in a humid chamber in RT. The slides were than washed and incubated with AB Solution (VECTASTAIN Elite ABC Hrp Kit, vector laboratories, PK6100) for 1 hour. Next, slides were washed again in PBS and incubated with DAB substrate (SIGMAFAST™ 3,3′-Diaminobenzidine tablets, Sigma D4293). Finally, the slides were washed again in PBS and covered with DPX mountant for histology (Sigma 4451). For IF staining: on the following day, slides were washed with PBS, and incubated with the appropriate fluorescently labelled secondary antibodies for 1.5 hours in a humid chamber in RT. Finally, slides were vertically air-dried and covered with DAPI flouromount-G (Southern biotech 0100-20). For BrdU staining, mice were injected ip with BrdU (Sigma B5002) 21 days prior to brain extraction.

### Antibodies

IHC and IF: Rabbit anti POGZ (ab171934); Mouse anti NeuN (MAB377); Mouse anti Anti-CUTL1/CUX1 (ab54583); Rat anti CTIP2 (ab18465); Rabbit anti TBR1 (Merck AB10554); Rabbit anti Ki67 (ab15580); Rat anti BrdU (ab6326) Mouse anti DCX (sc271390); Mouse anti SOX2 (sc17320); Goat anti NeuroD (sc1804); Rabbit anti PROX1 (Merck AB5475); Rabbit anti PAX6 (Merck AB2237); Mouse anti CNPase (ab6319); Mouse anti Calbindin (Merck C9848); Rabbit anti GFAP (ab7260); goat anti rabbit Alexa fluor 568 (Invitrogen A-11011); goat anti mouse Alexa fluor 488 (Invitrogen A-11001). WB Primary antibodies: Rabbit anti POGZ (ab167408) 1:500; Rat anti βTubulin (ab6160) 1:10000. WB Secondary abs: Donkey anti Rabbit IgG HRP (ab7083); Donkey anti Rat IgG HRP (ab102265).

### Golgi staining and analysis

Golgi staining was done using the “super Golgi kit” (Bioenno 003010). 2-month-old mice were euthanized using Isoflurane USP (Piramal critical care, NDC 60307-110-25). Their brains were then removed and immersed in 10ml of impregnation solution for 12 days. The rest of the protocol was done according to manufacturer instructions. Dendritic spines in the dentate gyrus were counted at 40μm distance from the soma and on a 30μm dendritic length.

### Behavioral assays

Mice were group-housed in a room with a 12-h-light, 12-h-dark cycle (lights on at 7:00) and with access to food and water *ad libitum*. All tests were conducted during the light cycle. Behavioral assays were performed with male and female mice at 8-16 weeks of age between 9:00 and 18:00, unless indicated otherwise. Each apparatus was cleaned with diluted EtOH solution before testing of each animal to prevent bias due to olfactory cues. Before all assays, mice were habituated to the testing facility for at least three days, and specifically to the testing room for 1-2 hours prior to the experiment. The behavioral assays were conducted blinded to genotypes during testing and analysis.

#### Three-chamber social interaction test

The three-chamber social interaction apparatus is divided into three interconnected compartments (35X20 each). Each side contains a wired cylinder for interaction (referred to as “cup”; 10.5cm height × 10cm bottom diameter). Test mouse was habituated to the apparatus for 5 minutes. Then, an unfamiliar mouse (Stranger 1) was randomly placed in the cup in one of the side compartments, while the other cup remained empty. The test mouse was re-introduced to the apparatus and allowed to explore all three chambers for a 10-minutes session. Subsequently, a novel unfamiliar mouse (Stranger 2) was placed in the previously empty cup and the test mouse had another 10 minutes to explore the apparatus. The test sessions were recorded using EthoVision XT11 tracking system (Noldus, Leesburg, VA). Both sessions were done in a dark room.

#### Marble burying test

Mice were habituated to a novel testing cage (35×20×18 cm^2^) containing a 5-cm layer of chipped cedar wood bedding for 10 minutes. The mice were returned to their home cage while 20 glass marbles were aligned equidistantly 4 x 5 in the testing cage. Mice were then introduced again to the testing cage and given 30 minutes to explore it. At the end of the session, the number of buried marbles was recorded. A marble was considered buried if more than half of it was covered with bedding. Results were calculated and plotted as the percentage of marbles buried for each genotype.

#### Open Field

The Open Field environment, consisted of a white circular plexigel arena (55cm diameter) was divided into “center” (28cm diameter) and “periphery” (the outer part of the circle). Mice were placed in one of four edges of the arena (N, W, S, and E) randomly, and their free movement was recorded for 5 minutes using the EthoVision XT11 tracking system (Noldus, Leesburg, VA).

#### Elevated Plus Maze

The Elevated plus maze apparatus consisted of four elevated arms (52cm from the floor). The arms were arranged in a cross-like disposition, with two opposite arms being enclosed (30×5×25 cm^3^) and two being open (30×5 cm^2^), having at their intersection a central square platform which gave access to any of the four arms. All floor surfaces were covered with yellow tape to create a high contrast for the tracking system. Mice were placed in the central platform facing an open arm and could explore the apparatus for 5 minutes. The session was recorded using EthoVision XT11 tracking system (Noldus, Leesburg, VA)

#### Buried food olfactory test

Mice were food deprived for 24 hours prior to the testing day. Test mouse was habituated to a novel testing cage (35×20×18 cm^2^) containing a 3cm layer of chipped cedar wood bedding for 5 minutes. Then, the mouse was removed, and a small cheese cube (1.5 gr) was hidden in one corner of the cage under the bedding. During the test, each mouse was placed into the center of the testing cage, and the latency to find the cheese was measured.

#### Social odors habituation / dishabituation

Prior to testing, experimenters prepared triplicates of cotton swabs soaked with mice urine and triplicates of cotton swabs soaked with a smell of a dirty cage. The odor samples were age- and sex-matched to the study mice. Test mouse was habituated to a new, clean testing cage (35×20×18 cm^2^) containing a clean odorless cotton swab for 30 minutes. During the experiment, each odorous swab was lowered to the cage for 2 minutes and the mouse was free to sniff it. The order of samples was: urine1, urine2, urine3, cage1, cage2, cage3 with a 1-minute interval between them. In all trials, the time the mouse spent sniffing the swab was recorded using a manual timer.

#### Rotarod

An accelerating Rotarod (Ugo Basile, Cmoerio VA, Italy) was used in this experiment. Five mice were placed simultaneously on the stationary rod (3cm diameter). Once all mice were stable on the rod, the motor was turned on and the rod rotation was continuously accelerated (from 4 to 40 rpm over 5 minutes). Each mouse was given three successive trials (30 minutes interval between trials) for a maximum of 7 minutes per trial. Results were measured as the time the mouse stayed on the accelerating rod before falling.

#### Horizontal bars

Motor coordination and forelimb strength test was performed as previously described^55^. Briefly, the bars were made of steel, 38cm long and held 49cm above the bench surface by a support column at each end. Two bar diameters were used for the experiment, 2 and 4-mm. Test mouse was held by the tail and raised to grasp the horizontal bar in the middle with the forepaws only. The criterion point was either a fall from the bar, or the time until one forepaw touches the support column. Maximum test time was 30 seconds. Mice were first tested on the 2-mm bar and then the 4-mm bar after 1-2 minutes resting period. Performance (Time until falling or touching the support column) scoring was as follows: 1-5 sec = 1; 6-10 sec = 2; 11-20 sec = 3; 21-30 sec = 4; after 30 seconds or placing a forepaw on the side column = 5. Final score was calculated as the sum of scores from both bars tests.

### Gene expression analysis

Adult mice hippocampi and cerebella (three males and three females from each genotype) were dissected and RNA was extracted using Qiagen RNeasy lipid tissue mini kit (Qiagen 74804). The RNA integrity was measured using the Agilent BioAnalyzer machine (RNA integrity number (RIN) was above 8.5). Libraries of mRNA for sequencing were made by pulldown of poly (A) RNA using Truseq RNA sample preparation kit (illumina). Library quantity and pooling were measured by Qubit (dsDNS HS) and quality was measured by tape station (HS). Sequencing was done using Next seq 500 high output kit V2 75 cycles (illumina) on illumina’s NextSeq 500 system with 43bp paired-end reads (resulting 50-70 million reads per sample). Samples denaturing and loading was according to manufacturer instructions.

Reads from RNA-Seq were aligned to the mouse genome (mm9), and aligned reads were counted at the gene level using STAR aligner (v250a). Reads that mapped to exons 13-19 at the *Pogz* gene were used to verify the genotypes. Raw count data was used for differential expression analysis using edgeR ^56^. Genes with counts per million (CPM) above 1 in at least six individual samples were included in the analysis. The differential expression analysis was performed using a generalized linear model that included sex and genotypes. Differences were considered statistically significant for FDR < 0.05 (calculated by edgeR package). Exact binomial test was used to test for the unequal proportion of upregulated and downregulated genes. Enrichment of gene ontology was analyzed using the Enrichr online tool ^34,35^.

### Slice Preparation for Electrophysiology recordings

Mice (8-12 weeks old) were anesthetized with pentobarbitone (60 mg/kg) and the brain was removed. The cerebellum was rapidly cut and placed in ice-cold physiological cutting solution containing the following (in mM): 124 NaCl, 2.4 KCl, 1 MgCl_2_, 1.3 NaH_2_PO_4_, 26 NaHCO_3_, 10 glucose, 2 CaCl_2_, saturated with 95% O_2_/5% CO_2_, pH 7.4 at room temperature. Parasagittal slices (250 μm) of the vermal and hemispherical area were cut using a microslicer (7000 SMZ, Campden Instruments, UK). The slices were incubated with oxygenated physiological solution and maintained at 34 degrees. After 1 hour incubation, the slices were transferred to a recording chamber and maintained at room temperature under continuous superfusion with the oxygenated physiological solution.

### Whole Cell Recordings

Neurons were visualized using differential interference contrast, infrared microscopy (BX61WI, Olympus, Tokyo, Japan). Purkinje cells were identified by the location of their somata between the granular and molecular layers, by soma size and by the presence of a clear primary dendrite. All recordings were performed at room temperature. For current clamp experiments, bath solution was the same as the cutting solution. Borosilicate pipettes (4-6 MΩ, pulled on a Narishige pp-83 puller.) were filled with internal solution containing the following (in mM): 145 potassium-gluconate, 2.8 KCl, 2 NaCl, 0.1 CaCl_2_, 2.4 Mg-ATP, 0.4 GTP, 10 HEPES and 1 EGTA (pH adjusted to 7.3 with KOH). Signals were recorded at 10 kHz and low-pass filtered at 3 kHz. Electrode access resistance was routinely checked, and recordings with values larger than 20 MΩ were not included in the analysis. Experiments were discarded if more than 20% change in holding current was needed to maintain the resting potential. AP threshold was calculated from phase plot (dV/dt) analysis of a single AP plotted versus membrane voltage. The point in the phase plot where the increase in dV/dt was larger than 10 mV/ms was defined as the AP threshold. Input resistance was calculated by fitting the voltage response during the short current injection to a double exponential function and the difference between the estimated steady state value and the initial membrane voltage was divided by the current injection amplitude. For I-f protocol PCs were held at approximately −70 mV, and 1 sec current pulse was given every 12 sec. In voltage clamp experiments, bath solution contained the following (in mM): 124 NaCl, 5 KCl, 2 CaCl2, 1.3 NaH_2_PO_4_, 1 MgCl_2_, 10 glucose, 26 NaHCO_3_, 1 NiCl_2_, 0.1 CdCl_2_, 1 BaCl_2_, 5 TEACl, 0.0005 TTX, 1 4-aminopyridine (4-AP). Borosilicate pipettes were filled with the same internal solution as in current clamp experiments. PCs were held at −50 mV, and 4 sec voltage pulses were given, every 30 sec, from −10 to −40 mV and 2 sec pulses from −50 to −70 mV, all in 10-mV steps. Whole cell capacitive transients and leak currents were not compensated during these recordings. Data were sampled at 10 kHz as in current-clamp experiments, and low-pass filtered at 1 kHz. Voltage clamp recordings were commenced 5-10 min after seal breaking. For 4 sec pulses, I_h_ current was defined as the difference between the minimal current recorded immediately after the ‘leak’ current and the current measured at the end of the voltage step. As for the 2 sec pulses, since the current has not reached steady-state, we fitted the current changes during the voltage step by either 1 or 2 exponents, and the estimated steady state value was subtracted from the minimal current recorded immediately after the ‘leak’ current. Analysis of I_h_ kinetics was done only on traces that were successfully fitted with 2 exponents, which were most of the traces. A dedicated software based on the LabView platform (National Instruments) was used for monitoring and data acquisition.

### In-vivo extracellular PC recordings

Mice were anesthetized with an I.P. Injection of a mixture containing ketamine (100 mg kg−1) and medetomidine (10 mg kg−1). Depth of anesthesia was sufficient to eliminate pinch withdrawal. Anesthesia was maintained throughout the experiment by I.P. Injection of additional supplements of ketamine (25 mg kg−1) administered as necessary (typically every 0.5–;1 h). Body temperature was kept at 37 degrees using a heating blanket. Craniotomy was performed 6.0 mm caudal to Bregma and between midline and 1.5 mm lateral from midline. Recordings were obtained using borosilicate pipettes (3-5MΩ) pulled on a Narishige pp-83 puller, filled with artificial cerebrospinal fluid (ACSF) containing: NaCl 125 mM, KCl 2.5 mM, NaHCO_3_ 25 mM, NaH_2_PO_4_ 1.25 mM, MgCl_2_ 1 mM, glucose 25 mM and CaCl_2_ 2 mM. Experiments were conducted only on male mice.

### Statistical analysis

Analysis and generation of plots were performed using custom scripts in R. Statistical testing was performed using linear models that included the number of intact *Pogz* alleles, the animal sex, and the interaction between them as the explanatory variables. In addition, a Tukey’s honest significance test was used to test for the pairwise differences between genotypes. For comparisons across time points, a model that included repeated-measures was used (e.g. rotarod and olfactory assays).

## Acknowledgments

This research was supported by the Israel Science Foundation (grant no. 940/13 and 575/17 to SS) and funded in part with Federal funds from the Frederick National Laboratory for Cancer Research, NIH, under Contract HHSN261200800001E (to JRK). The content of this publication does not necessarily reflect the views or policies of the Department of Health and Human Services, nor does mention of trade names, commercial products or organizations imply endorsements by the US Government.

